# A yeast-based system to study SARS-CoV-2 M^pro^ structure and to identify nirmatrelvir resistant mutations

**DOI:** 10.1101/2022.08.06.503039

**Authors:** Jin Ou, Eric M. Lewandowski, Yanmei Hu, Austin A. Lipinski, Ryan T. Morgan, Lian M.C. Jacobs, Xiujun Zhang, Melissa J. Bikowitz, Paul Langlais, Haozhou Tan, Jun Wang, Yu Chen, John S. Choy

## Abstract

The SARS-CoV-2 main protease (M^pro^) is a major therapeutic target. The M^pro^ inhibitor, nirmatrelvir, is the antiviral component of Paxlovid, an orally available treatment for COVID-19. As M^pro^ inhibitor use increases, drug resistant mutations will likely emerge. We have established a non-pathogenic system, in which yeast growth serves as a proxy for M^pro^ activity, enabling rapid identification of mutants with altered enzymatic activity and drug sensitivity. The E166 residue is known to be a potential hot spot for drug resistance and yeast assays showed that an E166R substitution conferred strong nirmatrelvir resistance while an E166N mutation compromised activity. On the other hand, N142A and P132H mutations caused little to no change in drug response and activity. Standard enzymatic assays confirmed the yeast results. In turn, we solved the structures of M^pro^ E166R, and M^pro^ E166N, providing insights into how arginine may drive drug resistance while asparagine leads to reduced activity. The work presented here will help characterize novel resistant variants of M^pro^ that may arise as M^pro^ antivirals become more widely used.

## Introduction

The evolution of new SARS-CoV-2 variants that evade vaccines, cause breakthrough COVID-19 infections in vaccinated individuals, and the limited vaccine availability in many parts of the world, highlight the need for complementary approaches^1^. Antiviral drugs provide an important alternative and can contribute to minimizing disease severity and death. The SARS-CoV-2 main or 3C-like protease (M^pro^ or 3CL^pro^) is essential for viral replication and is a promising drug target ^2,3^. There have been intense efforts to repurpose or to develop new drugs that directly target M^pro 4,5^. In December 2021, emergency authorization use of Paxlovid to treat COVID-19 was granted by the US Food and Drug Administration ^6^. Paxlovid is a combination of the M^pro^ inhibitor, nirmatrelvir, and the cytochrome CYP3A inhibitor, ritonavir, which slows metabolism of nirmatrelvir ^7,8^. Currently, there are several other M^pro^ inhibitors in clinical trials, including PF-07304814, the phosphate form of PF-00835231^9,10^. As M^pro^ inhibitors become more widely used the emergence of resistant mutations will increase as greater selection pressure is present in the population.

Knowledge of resistant mutants can inform on drug design modifications to identify new drugs that target resistant variants. However, standard approaches to characterize resistant mutants using live virus^11^, recombinant proteins, and in vitro assays can be highly limiting due to infrastructure requirements, cost, and time^12^. Here we report a yeast system that is non-pathogenic, rapid, inexpensive, and reports on M^pro^ activity and drug resistance simply by measuring yeast growth. Using this assay, we found that compared to wild-type, the E166R mutation conferred strong nirmatrelvir resistance (*K*_*i*_ > 1000-fold). As the E166 site appears to be a hot spot for drug resistance from *in vitro* viral evolution experiments^13,14^, we solved the structures of two substitution mutants M^pro^ E166N and M^pro^ E166R, revealing how E166 mutations may compromise activity versus drug resistance, respectively. Our results demonstrate the yeast system can be a reliable tool to determine the activity and drug responses of M^pro^ mutants. Results from the yeast assays can help rapidly prioritize mutants for further analysis using more resource intensive systems. In doing, so we can efficiently test M^pro^ mutants as they arise in the population and aid in mitigating COVID-19 infections.

## Results

### SARS-CoV-2 M^pro^, PL^pro^, spike, and helicase proteins are toxic in *S. cerevisiae*

We expressed six SARS-CoV-2 (Wuhan-Hu-1) NSPs and the structural genes, spike, M, E and N^15^ to determine if any would result in growth effects (Fig. 1A and Fig. S1A). We observed no marked growth phenotypes as determined by spot tests when M, E, N, NSP7, NSP8, or NSP12 were expressed (Fig. S1B). In contrast, spot tests revealed nearly a complete absence of growth when cells expressed NSP3 (PL^pro^), NSP5 (M^pro^ or 3CL^pro^), NSP13 (Helicase), and spike (Fig. 1A and 1B). Analysis of growth profiles of cells expressing PL^pro^, M^pro^, Helicase, and spike showed all four genes caused a reduction in growth. M^pro^ and the Helicase were the most toxic conferring a ∼70 to 80% reduction in total growth by 72 hours compared to cells carrying empty vector (Fig. 1A). As M^pro^ is highly conserved between classes of coronavirus and a key drug target we focused our efforts on using the yeast system to study M^pro^ structure and function.

**Fig. 1.**
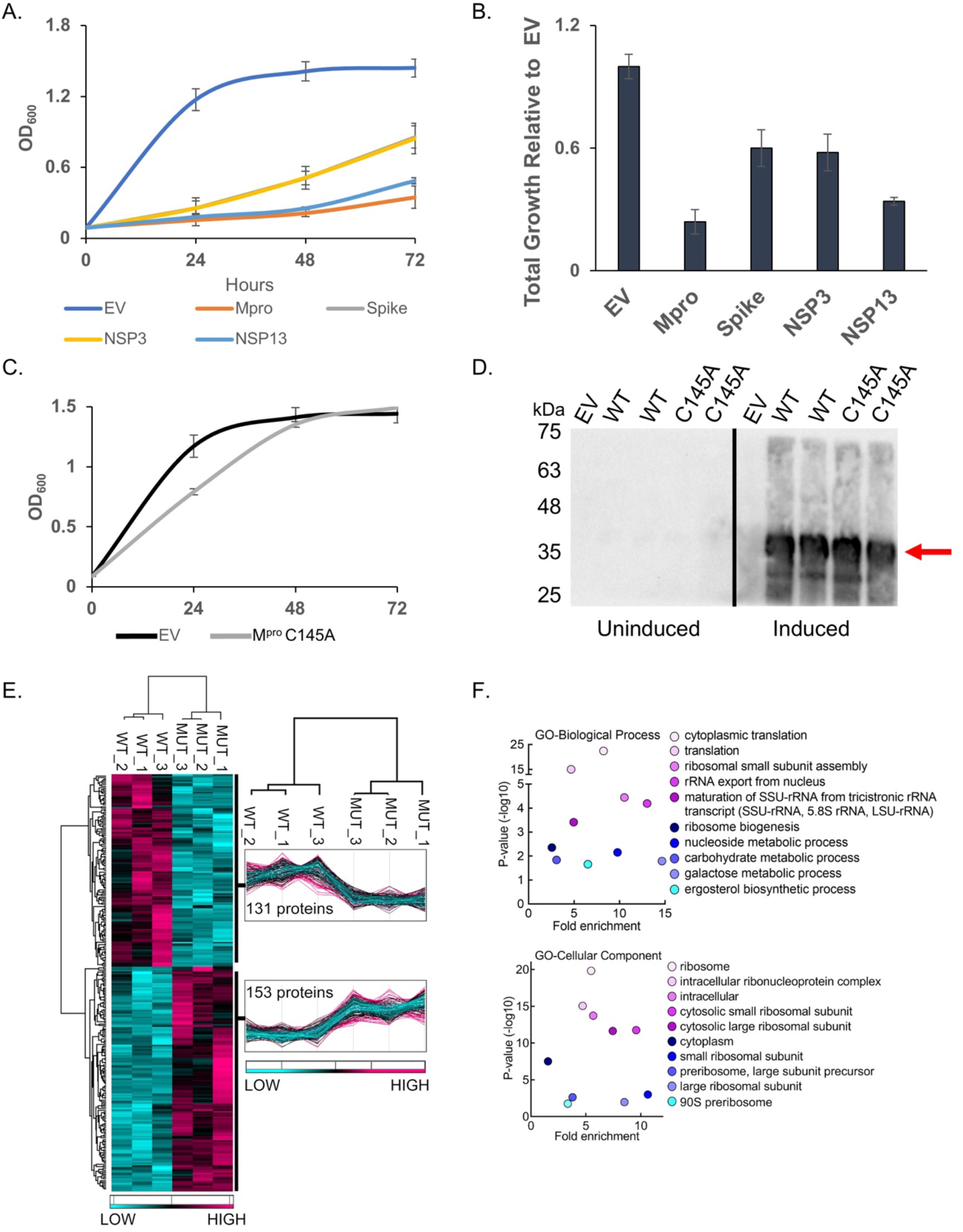
M^pro^ confers a significant reduction in growth in yeast caused by decreases in a variety of cellular proteins. A) The indicated SARS-CoV-2 genes under a galactose inducible promoter were expressed in yeast and conferred growth defects compared to empty vector (EV). B) Bar graph shows the total growth of cultures after 72 hours normalized to EV. C) Expression of the catalytically inactive M^pro^ C145A mutant does not confer a growth reduction and yeast grow similarly to EV control cells. D) Protein levels of the M^pro^ C145A mutant and wild-type M^pro^ (WT) are comparable. Shown are two biological replicates for each form of M^pro^. E) Total protein lysates made from yeast expressing the wild-type M^pro^ (WT) or M^pro^ C145A mutant (MUT) were subjected to mass spectrometric analyses revealing 153 proteins were higher in abundance in the mutant relative to the wild-type. F) Gene Ontology (GO) analyses indicates an enrichment of proteins with functions in translation that are significantly reduced in the presence of M^pro^ versus M^pro^ C145A. Plots in A and B show averages from three biological replicates and error bars are standard deviations.

### Growth defect conferred by M^pro^ expression depends on its catalytic activity and associated with decreased abundance in essential and non-essential yeast proteins

To determine if the growth reduction depended on M^pro^ proteolytic activity we constructed a catalytic mutant of M^pro^ by replacing the key cysteine at position 145 to an alanine, which prevents the initial protonation step needed for peptide bond hydrolysis^16,17^. Liquid growth assays showed that yeast expressing the M^pro^ C145A mutant grew as well as the yeast control carrying empty vector (Fig. 1C). Western analysis showed that yeast expressed similar levels of wild-type and M^pro^ C145A mutant (Fig. 1D). These results demonstrate that the growth reduction observed in yeast expressing M^pro^ is dependent on its proteolytic activity.

Next, we measured the relative abundance of proteins in yeast expressing M^pro^ compared to yeast expressing the M^pro^ C145A catalytic mutant to determine the mechanism(s) that lead to loss of cell viability. Whole cell lysates were made from three independent cultures of cells expressing wild-type M^pro^ or the catalytic M^pro^ C145A mutant (Fig. 1E and Fig. S2). The biological replicates were highly reproducible, and we observed peptides from 153 proteins were significantly reduced in yeast expressing M^pro^ compared to the M^pro^ C145A mutant (Fig. 1E.) Gene ontology analysis revealed an enrichment for genes with functions in translation (Fig. 1F). In particular, multiple ribosomal proteins and translational regulators were reduced. There were a number of proteins that were significantly enriched in the M^pro^ catalytic mutant with functions in a variety of activities beyond translation and several are known to be essential. These results show that expression of M^pro^ leads to decreases in a variety of proteins and eventual loss of translation that is likely the cause of the growth defects.

### Nirmatrelvir restores growth to yeast expressing M^pro^ from multiple coronaviruses

Considering that the growth reduction conferred by M^pro^ activity is dependent on its proteolytic activity we tested if treating yeast with nirmatrelvir, would suppress the growth reduction. We tested nirmatrelvir at several concentrations and observed no cytotoxic effects (Fig. S3A). Treating cells with increasing doses of nirmatrelvir led to a corresponding increase in growth (Fig. 2A and Fig. S2A). At 100μM and 200μM of nirmatrelvir, growth was restored to similar levels as cells carrying empty vector (Fig. 2A). As a metric to compare the effects of nirmatrelvir, we estimated the concentration of drug required to restore 50% of growth (RC_50_) relative to that of untreated M^pro^ expressing cells. Based on this criterion we calculated RC_50_ for nirmatrelvir to be 110.47 ± 4.76μM (Fig. 2A and 2F). To determine if M^pro^ from other coronaviruses could be studied similarly, we tested the recent Omicron variant, M^pro^ P132H, which is currently the dominant form of M^pro^, and M^pro^ from SARS-CoV-1 and Bat-CoV-HKU9. We observed that in all cases M^pro^ conferred a significant growth reduction (Fig. 2B and Fig. S4A). Nirmatrelvir has been reported to have broad M^pro^ specificity^7,9^. Consistent with this work, we observed that nirmatrelvir could restore growth in yeast expressing M^pro^ from all three forms of M^pro^ (Fig. 2B and Fig. S4B).

**Fig. 2.**
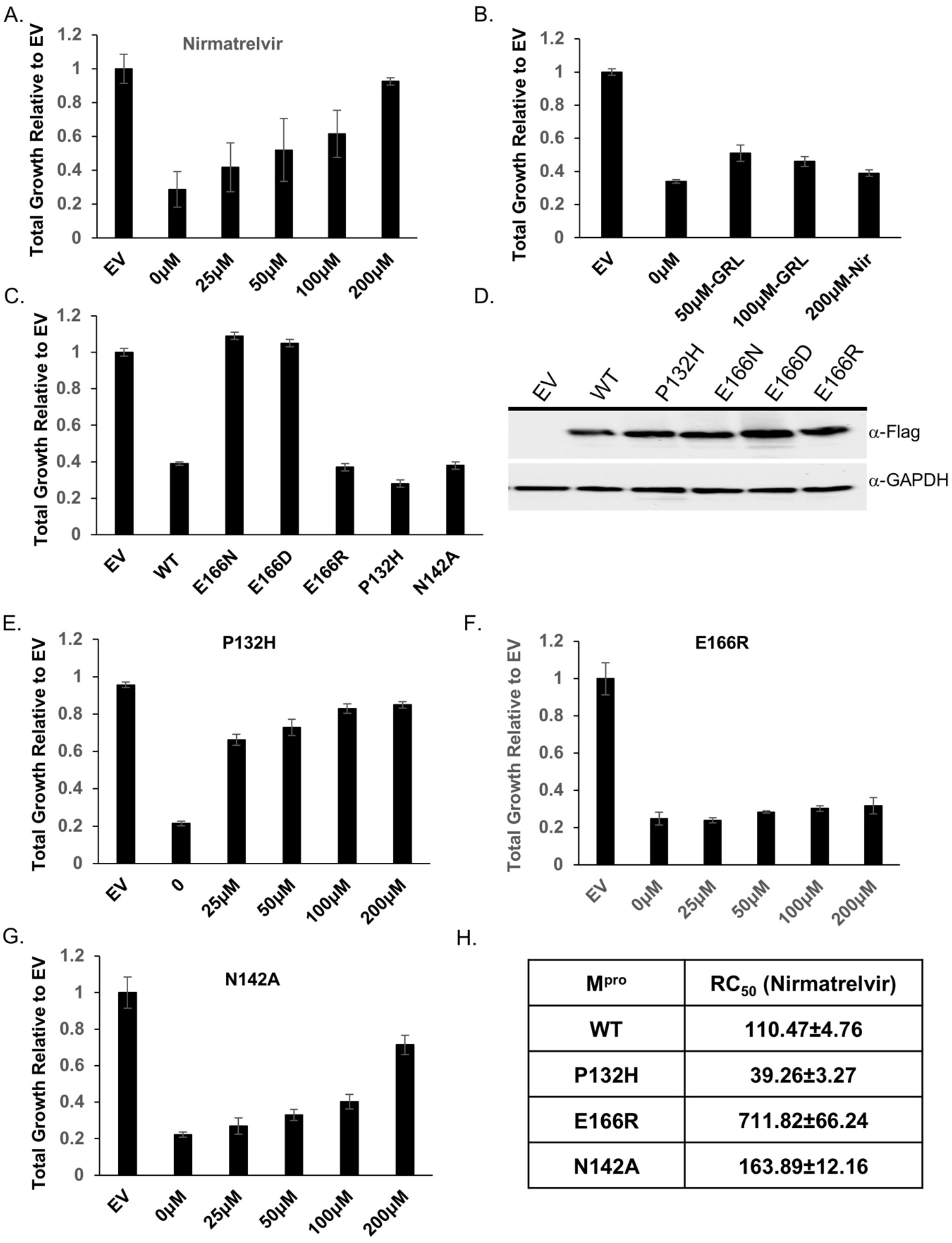
Yeast growth assays identify nirmatrelvir resistant M^pro^ mutants. A) Total growth of cultures after 72 hours expressing M^pro^ in the presence of increasing doses of nirmatrelvir normalized to growth of yeast carrying empty vector (EV) are plotted. Nirmatrelvir is effective at protecting yeast from the toxicity of M^pro^ as growth is restored to levels comparable to EV. B) Yeast expressing PL^pro^ show a growth reduction and treatment with nirmatrelvir (nir) does not rescue the growth defect. On the other hand, treating cells with GRL0617 (GRL) restores some growth. C) Yeast expressing substitutions E166D and E166N grow as well as EV but E166R, P132H, and N142A results in significant growth reduction comparable to wild-type M^pro^. D) Western analysis shows that mutants and wild-type M^pro^ are expressed at comparable levels. E - G) Cells expressing P132H and N142A remain sensitive to nirmatrelvir, indicated by growth recovery, but E166R appears to be resistant as there is a lack of growth even when treated with 200µM of nirmatrelvir. H) RC_50_ measurements of each mutant in response to nirmatrelvir treatment. For all experiments, at least three biological and three technical replicates were performed for EV, wild-type and mutant M^pro^. Error bars represent standard deviations.

To determine the specificity of the restored growth conferred by nirmatrelvir in cells expressing M^pro^, we tested the effects of nirmatrelvir on cells expressing PL^pro^. The growth reduction associated with expression of the PL^pro^ (Fig. 1A) should not be inhibited by this drug. We treated cells expressing PL^pro^ with 200µM of nirmatrelvir and observed no improvement in growth (Fig. 2B). On the other hand, 50 µM and 100µM of GRL0617, an inhibitor of PL^pro 18^ was associated with partial recovery of growth (Fig. 2B). Together, these observations show that the restoration of growth conferred by nirmatrelvir in cells expressing M^pro^ is specific to M^pro^ rather than a non-specific effect on yeast physiology.

### Characterization of potential nirmatrelvir resistant mutations in M^pro^

We tested if growth of yeast expressing M^pro^ could be used as a proxy for M^pro^ activity. Thus, providing a system to rapidly determine structure function relationships as they relate to activity and drug resistance. A variety of interactions (H-bonds, salt-bridges, van der waals) mediate binding between the catalytic site of M^pro^ and inhibitors ^19-21^. While knowledge of the residues in contact with the inhibitor can inform predictions that may compromise inhibitor binding it is not obvious what amino acid substitutions would maintain M^pro^ activity toward substrate while compromising inhibitor interactions. With our yeast system we can easily test the effect of substitution mutations and rapidly determine if the mutations alter catalytic activity and sensitivity to inhibitor(s) by following growth phenotypes. To determine the feasibility of this approach we focused on E166, and N142 as these two residues form direct interactions with inhibitors and substrates ^19,22^.

We tested substitutions of E166 with three different amino acids that are yet to be dominant or present in the population. The following mutants predicted to be conserved (E166D), as the negative charge is maintained but with one less carbon in the side-chain; non-conserved (E166N), as asparagine is uncharged and has one less side chain carbon; and another non-conserved (E166R) substitution in which the arginine side chain is longer and positively charged were tested. We observed that all three substitutions were expressed at the same levels as wild-type M^pro^ but M^pro^ E166D and M^pro^ E166N mutants did not cause a reduction in growth and grew as well as empty vector controls (Fig. 2C and 2D, Fig. S5A). These results indicate that M^pro^ E166D and M^pro^ E166N may have defects in their enzymatic activities. However, the M^pro^ E166R mutant conferred a growth reduction that matched the wild-type M^pro^, suggesting that its catalytic activity was intact (Fig. 2C and 2D, Fig. S5A).

Next, we challenged cells expressing M^pro^ E166R with increasing concentrations of nirmatrelvir (25μM, 50μM, 100μM, or 200μM) and observed no significant improvement in growth remaining nearly identical to the untreated culture of M^pro^ E166R expressing cells (Fig. 2E and Fig. S5b). Based on these experiments, the RC_50_ for nirmatrelvir is 711.82 ± 66.24μM, a ∼7-fold increase in RC_50_ compared to wild-type M^pro^. These results suggest that the E166R mutation confers resistance to nirmatrelvir.

We constructed a substitution at position N142, which is known to contribute to inhibitor and substrate binding^7^ and is yet to be present in the population. To inform on the specific substitution to make we used a distantly related M^pro^ from the gamma-coronavirus, IBV, which is conserved but displays slight divergence from SARS-CoV-2 M^pro9^. We replaced N142 with alanine (M^pro^ N142A), as alanine is found in the IBV M^pro^ at the homologous site^23^. We observed a reduction in growth comparable to M^pro^ when M^pro^ N142A was expressed (Fig. 2C) showing that it remained active. The RC_50_ for nirmatrelvir increased modestly by ∼1.5-fold (Fig. 2C). These results show that the substitution mutant E166R leads to nirmatrelvir resistance while N142A results in little difference from wild-type and E166N and E166D cause a loss in activity.

### *In vitro* protease assays confirm that MproE166R is highly resistant to nirmatrelvir

The results from the yeast assays suggest Mpro E166R confers resistance to nirmatrelvir (∼7-fold increase in RC50 vs WT). To determine how well yeast growth assays correlated with standard enzymatic assays we directly measured protease activity using recombinant Mpro, Mpro E166N, Mpro E166R, and Mpro N142A. First, we measured the catalytic efficiencies for all four forms of Mpro (Fig. 3A). Compared to wild-type Mpro, the catalytic efficiencies (kcat/Km) of Mpro E166R was decreased by ∼16-fold, while Mpro N142A displayed a slight increase of 1.4-fold. In contrast, E166N was nearly inactive with *k*_*cat*_*/K*_*m*_ of 132 S^-1^M^-1^, a 83.5-fold reduction compared to WT. The enzymatic assay results confirmed that the lack of toxicity of E166N in the yeast growth assay was due to the loss of catalytic activity (Fig. 2C). To determine the response of the mutants (M^pro^ E166R, and M^pro^ N142A) to inhibitors compared to wild-type, we measured the IC_50_ and *K*_*i*_ for nirmatrelvir, and two other M^pro^ inhibitors PF-00835231, and GC-376^9,24^. We observed for M^pro^ E166R, increases in IC_50_’s of ∼143-fold for nirmatrelvir, ∼52-fold for PF-0083521, and ∼52-fold for GC-376. On the other hand, M^pro^ N142A, only minor increases in IC_50_’s of ∼1.4-fold for nirmatrelvir, ∼1.9-fold for PF-0083521, ∼1.1-fold for GC-376 (Fig. 3B). The *K*_*i*_ values for the inhibitors in assays with M^pro^ E166R were increased by ∼1620-fold for nirmatrelvir, ∼423-fold for PF-0085231, and ∼37-fold for GC-376 (Fig. 3C). Nearly no difference in *K*_*i*_ values from assays with M^pro^ N142A, ∼1.2-fold for nirmatrelvir, ∼0.9-fold for PF-0085231, ∼1.5-fold for GC-376) (Fig. 3C). The enzymatic assays confirm the results from the yeast assays showing that M^pro^ E166R is highly resistant to nirmatrelvir and also show that there is cross-resistance to PF-0085231 and GC-376 (Fig. 2D). Similarly, results from yeast assays of M^pro^ N142A mutant appears to correspond well to the *in vitro* assays as both show minor to no increases in resistance (Fig. 3). Furthermore, the M^pro^ E166N, which is not predicted to be catalytically active from the yeast assay, displayed >83-fold decrease in activity compared to wild-type in the *in vitro* assays. This result is completely consistent with observing no growth reduction when expressed in yeast. Taken together there is good correlation between the enzyme and yeast assays.

**Fig. 3.**
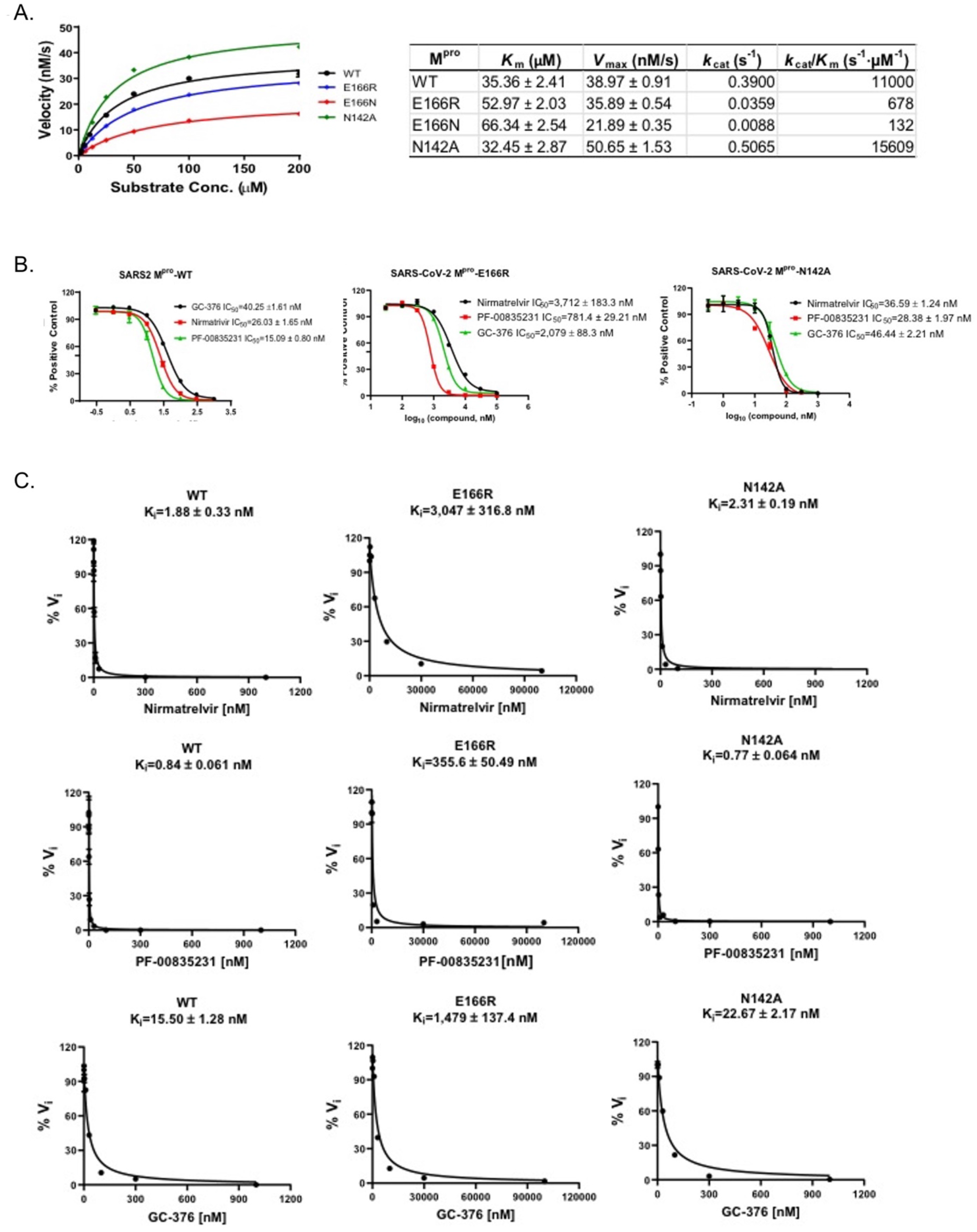
Enzymatic assays demonstrate that M^pro^ E166R is highly resistant to PF-00835231, nirmatrelvir, and GC-376. A) Michaelis–Menten plot of M^pro^ and its mutants with various concentrations of FRET substrate. The K_m_, V_max_, k_cat_, and *k*_*cat*_*/K*_*m*_ values are shown in the table on the right. B) The IC_50_ plots of nirmatrelvir, GC-376, and PF-00835231 against M^pro^, M^pro^ E166R, and M^pro^ N142A. C) K_i_ plots of nirmatrelvir, GC-376, and PF-00835231 against M^pro^, M^pro^ E166R, and M^pro^ N142A.

### Crystal Structure of Mpro^E166R^ reveals a loss of interactions leading to drug resistance

We were particularly interested in how replacing glutamate at position 166 with arginine led to a >1000-fold increase in resistance to nirmatrelvir while a substitution with asparagine led to an 83.5-fold decrease in enzymatic activity even though E166 is not known to be directly involved in catalysis. Toward addressing both questions, we solved the crystal structure of apo M^pro^ E166N and the complex structure of M^pro^ E166R with GC-376 at 2.3 and 2.1 Å resolution, respectively (Fig. 4). In the M^pro^ E166N mutant structure, N166 forms a hydrogen bond (HB) with H163, an interaction not observed between E166 and H163 in the wild-type M^pro^ structure (Fig. 4A). This new HB prevents H163 from hydrogen bonding with the glutamine side chain of the substrate, an interaction crucial to substrate binding. The binding of the substrate would therefore require N166 to adopt a different conformation, breaking the HB with H163 and increasing the energetic cost. These observations explain the drastic decrease of activity in the E166N mutant and lack of toxicity when expressed in yeast (Fig. 2C) bringing to light how residues outside of the catalytic core can influence substrate binding.

**Fig. 4.**
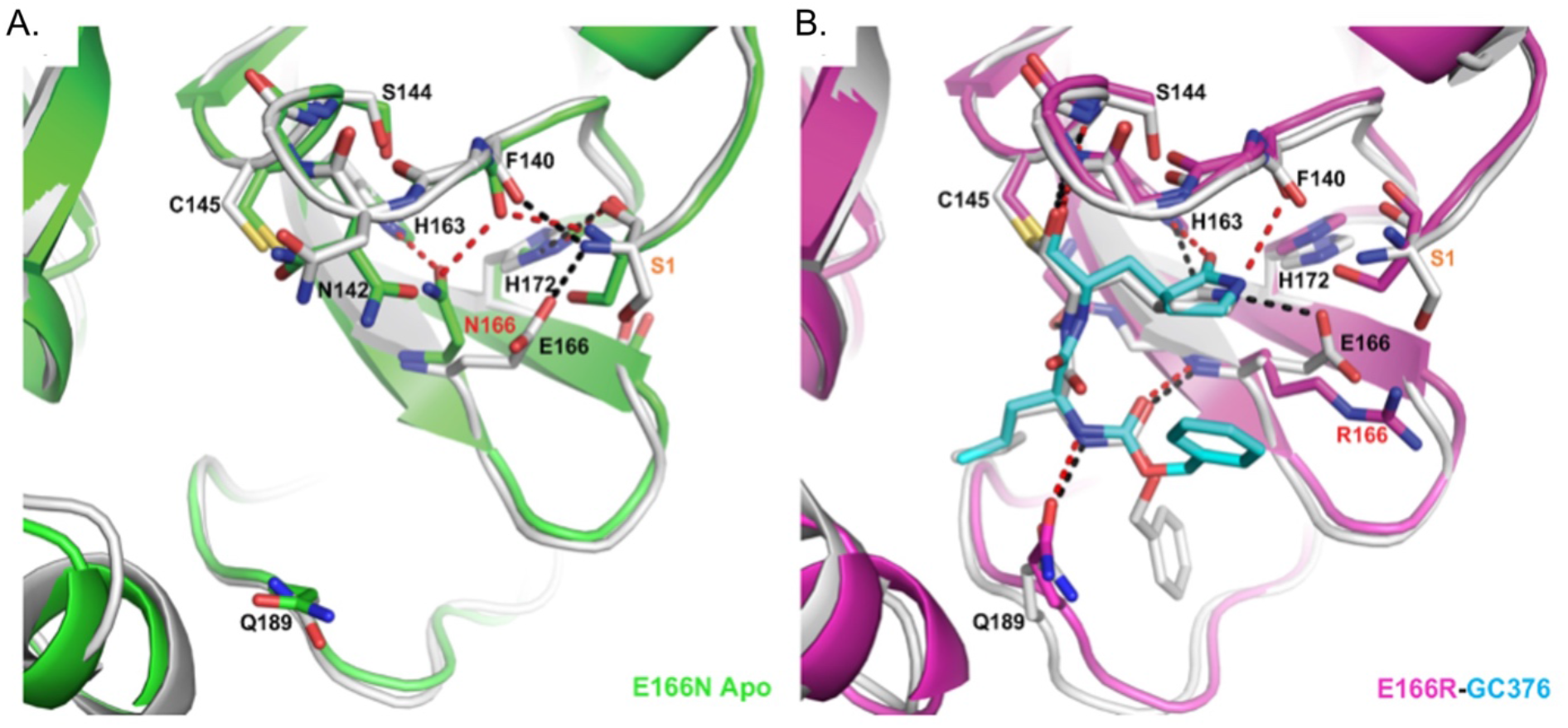
Crystal structures of E166R reveals structural basis for resistance and of E166N reveals basis for inactivity. A) Apo Mpro WT (white, PDB 7JP1) aligned with apo Mpro E166N (green, PDB 8DDI). B) Mpro WT GC376 complex (white, PDB 6WTT) aligned with Mpro E166R GC376 complex (magenta, PDB 8DDM). WT hydrogen bonds are shown as black dashes, and mutant hydrogen bonds are shown as red dashes. GC376 is shown in white for the WT structure and cyan for the mutant structure. Mutations are indicated with red text. Ser1 from an adjacent protomer is indicated with orange text.

In contrast, the longer and positively charged R166 side chain in the M^pro^ E166R mutant does not interact with H163, but rather extends into the solvent (Fig. 4B). Therefore, the S1 site is open for substrate binding. However, the E166R mutation does affect ligand binding in several aspects. The negatively charged E166 side chain forms two crucial HBs, one with the N-terminus of the neighboring M^pro^ protomer in the biological dimer, and the other with the pyrrolidone side chain of inhibitors (in both nirmatrelvir and GC-376) or with the glutamine side chain of the substrate as described above. The E166R mutation would abolish this direct HB with the substrate or inhibitor, resulting in the pyrrolidone ring of GC-376 forming an alternative weak HB with F140 (3.1 Å in length) in the mutant complex structure (Fig. 4B). In addition, the N-terminus of the enzyme interacts with both E166 and the backbone carbonyl group of F140, and plays an important role in maintaining the structural stability of the enzyme active site. The E166R mutation eliminates the salt bridge with the N-terminus of the adjacent protomer, and further introduces electrostatic repulsion leading to small yet significant changes in the N-terminus conformation. Consequently, the distance between the N-terminal amine group and the F140 carbonyl group increased from 2.6 Å in the WT to 3.7 Å in the M^pro^ E166R mutant, diminishing the HB. This in turn may destabilize the loop that F140 resides on and also contains other important structural features involved in enzyme catalysis and ligand binding, including the backbone amide groups of Gly143 and Ser144 that form part of the oxyanion hole to stabilize the reaction transition state. This loop also contains the peptide bond between Leu141 and Asn142 that interacts with the two extra carbon atoms of the inhibitor pyrrolidone ring, but not with the substrate glutamine side chain. Destabilization of the region near F140 may increase the entropic cost of binding to the rigid pyrrolidone ring of nirmatrelvir and GC-376, more than the smaller and more flexible substrate glutamine side chain. For similar entropic reasons, the HB between the pyrrolidone ring and E166 might contribute more to inhibitor binding than that between the more flexible glutamine side chain and E166 (Fig. 4B). Consequently, the E166R mutation may have a stronger effect on binding to inhibitors such as nirmatrelvir versus substrate.

## Discussion

In sum, we demonstrate that using yeast growth as a proxy for M^pro^ activity can be a reliable indicator of the effects that mutations in M^pro^ can have on its activity and potential for drug resistance. Yeast assays indicated that an E166R mutation was resistant to nirmatrelvir and *in vitro* enzyme assays confirmed this observation, revealing a ∼1600-fold increase in resistance. Furthermore, the C145A catalytic mutant^2^ and E166N mutant did not cause a growth reduction in yeast and enzyme assays showed that the E166N substitution confers a dramatic ∼83-fold decrease in activity. In yeast assays the N142A mutant displayed minor differences in drug sensitivity compared to wild-type (RC_50_ ∼1.5-fold more than WT), which was confirmed by our *in vitro* enzyme assays. Similarly, the P132H mutant remained sensitive to nirmatrelvir based on our yeast assay, potentially even more sensitive with an RC_50_ ∼2.8-fold less than WT. This is consistent with previous reports showing that the P132H mutant remains sensitive to nirmatrelvir in *in vitro* enzyme assays ^25-28^. It appears that M^pro^ mutants (i.e. E166R) that have a decrease in catalytic efficiencies of up to 16-fold compared to WT are still able to confer a marked reduction in yeast growth. This is important as resistant mutants are likely to reduce protein fitness ^29,30^. However, the yeast assay is unable to detect enhanced M^pro^ activity (e.g., M^pro^ N142A), which we observed in *in vitro* assays. This may have been due to the relatively small increase (1.4-fold). However, the enhanced activity associated with N142A suggests that M^pro^ can evolve to be a more active enzyme. It is possible that mutants which enhance M^pro^ activity can improve protein fitness when combined with resistant mutants that on their own may have reduced activity^14^. The crystal structure of E166R with GC-376 revealed loss of key hydrogen bonds with the pyrrolidone ring of GC-376 which can explain the increase in resistance to nirmatrelvir containing the same functional group. On the other hand, the E166N mutant which could be considered a more conserved change than E166R decreased activity by ∼83-fold and did not confer a growth reduction in the yeast assays. In turn the crystal structure shows that the asparagine prevents substrate binding through a new hydrogen bond with H163, providing a mechanism to explain the significant reduction in activity. The additional mutants at E166 that are associated with *in vitro* viral evolution experiments along with what we show here highlight the importance of this site in playing a role in nirmatrelvir resistance. Our crystal structure illuminates a structural mechanism to help explain how substitutions at E166 can either lead to loss of activity versus gain of resistance.

While the drug doses used with yeast are in the micromolar versus nanomolar range that is more typical of *in vitro* enzymatic or viral assays, we observed good correlations between the yeast and enzymatic assays for nearly all of the mutants tested. The higher concentrations of drug may be needed even though we deleted the major efflux pump, Pdr5, as yeast harbor a range of efflux activities ^31^, or possibly differences in permeability as a result of lipid composition differences from human cells, as well as potential drug interactions with the yeast cell wall ^32^. Additional differences observed between the yeast and enzymatic assays may be a result of having multiple substrates in yeast, additional complexity of the cellular proteome, differences in pH, salt, and oxidation levels.

Taken together, these results demonstrate that a non-pathogenic, rapid, inexpensive and highly accessible yeast-based method can be used to characterize mutants for both their effects on M^pro^ activity and their responses to inhibitor compounds. There are reports using yeast as a tool to screen for M^pro^ inhibitors or perform mutational analysis ^33,34^. These systems incorporate M^pro^ reporters and modification of M^pro^ to carry a N-terminal serine. Our work shows that measuring the effects of M^pro^ (with a N-terminal methionine) on yeast growth (without any reporters) can be a rapid and inexpensive approach to determine consequences of M^pro^ mutations on activity and drug response. The qualitative results from the yeast assays can be an important tool to help prioritize mutants of interest before moving ahead to more demanding viral based experiments. As more inhibitors are used in the general population there will be increasing selection pressures for drug resistant mutations that will go beyond the current set of mutants that are potentially drug resistant^35,36^. The yeast system reported here promises to be an invaluable tool in helping to combat future drug resistant mutations to stem the tide of COVID-19 infections.

## Materials and Methods

### Strains, media, and chemicals

All yeast strains carried a *pdr5*::*G418* deletion in the BY4741 background (*MATa his3Δ1 leu2Δ0 met15Δ0 ura3Δ0*). Yeast were grown in liquid synthetic complete (SC) media (0.17% yeast nitrogen base, 0.5% ammonium sulfate, amino acid mix with appropriate drop out as noted, 2% glucose) or on solid SC media containing 2% agar at 30°C. Media and reagents for culturing yeast were from United States Biological (Salem, MA). M^pro^ and PL^pro^ inhibitors were from MedChemExpress (Monmouth Junction, NJ) and Selleck Chemicals (Houston, TX). All other chemicals were from Sigma Aldrich (St. Louis, MO) or VWR (Radnor, PA).

### Expression of SARS-CoV-2 genes in yeast and mutagenesis

The indicated SARS-CoV-2 genes were codon optimized for yeast, tagged at the 3’ with a 3X-Flag epitope, carried on high copy plasmids and genes were under the control of the Gal1 promoter (see Table S1). Site directed mutagenesis was performed using In-Fusion Cloning Kit (Takara). Primers used for mutagenesis can be found in Table S2 and purchased from IDT DNA Technologies (Coralville, IA). The M^pro^ gene was sequenced to confirm that mutations were incorporated successfully.

### Yeast Transformation

A single yeast colony was used to inoculate 5ml liquid YPD (1% yeast extract, 1% yeast bacto-peptone, 2% glucose) and grown overnight at 30°C. The next day cells were washed and resuspended in 1ml lithium acetate/TE solution (100 m*M* lithium acetate, 10 m*M* Tris-HCl, 1 m*M* EDTA, pH 7.5). Cells were aliquoted (60 μl) into microcentrifuge tubes, followed by the addition of denatured salmon sperm DNA (50μg), 0.2μg of plasmid, 1ml polyethylene glycol (PEG) lithium acetate solution (40% (w/v) PEG 4000, 100 m*M* lithium acetate, 10 m*M* Tris-HCl, 1 m*M* EDTA, pH 7.5), and incubated for 45min at 30°C. This was followed by a 20min incubation at 42° and chilled for 2min on ice. Cells were washed and resuspended in 100μl H_2_O and plated on selective SC agar plates, incubated for ∼3 days at 30°C.

### Protein extraction and western analysis

Cells were grown overnight in 5 ml SC-Ura, 2% raffinose at 30°C. The next day, fresh cultures were started with optical density OD_600_ of 0.5 in 20 ml SC-Ura, 2% galactose at 30° for 6 hrs. Cells were then harvested, frozen in liquid nitrogen, and stored at −80°. For total protein extract, trichloroacetic acid was performed as described previously ^37^ and protein concentration was determined by BCA protein assay kit (Thermo scientific). Protein samples were separated by 4– 12% gradient SDS-PAGE (GenScript) and blotted onto nitrocellulose or PVDF membranes. The following primary antibodies were used at 1:5000 dilution: anti-FLAG antibody (GenScript), and anti-GAPDH antibody (Proteintech). Secondary anti-mouse IgG HRP antibody was used at 1:7000 dilution (Promega). ChemiDoc (Bio-Rad) imaging system was used to detect chemiluminescence signals from blots.

### Cell growth assays and RC_50_ measurements

Cells were grown overnight in 5ml SC-Ura, 2% raffinose at 30°C. The next day, fresh cultures were started with an OD_600_ of 0.1 in SC-Ura, 2% galactose, with or without inhibitors and transferred to to 96-well plates, incubated at 30°C on a a rotary shaker. Three independent transformants were used to test each form of M^pro^. Each transformant was sampled three times for each assay. The plate was transferred to a Tecan Infinite 200 PRO plate reader, and OD_600_ measurements were taken at 0, 24, 48 and 72 hours, with 5 flashes per well. Excel (Microsoft) was used to analyze the raw data. As a measure of inhibitory activity of nirmatrelvir we calculated a Recovery Concentration (RC_50_). The slopes from the dose responses were calculated and used to estimate the concentration of inhibitor that improves growth to half-maximal relative to empty vector control after 72 hours of growth.

### Yeast Proteomics

Cells were grown overnight in 10ml SC-Ura + 2% Raffinose media at 30°C. The next day, fresh cultures were started with OD_600_ of 0.1 in 100ml SC-Ura + 2% galactose at 30°C for 6 hr. Cells were then harvested, frozen in liquid nitrogen, and stored at −80°. Protein extraction was performed as described previously^37^ and protein concentration was determined using Pierce BCA protein assay kit (Thermo scientific).

To determine changes in the proteome associated with expression of M^pro^ versus M^pro^ C145A, in-solution tryptic digestion was performed as described^38^ followed by desalting with a Pierce Peptide Desalting Spin Columns per the manufacturer’s protocol (ThermoFisher Scientific, cat no. 89852) and the peptides were dried by vacuum centrifugation. 600 ng of the final sample was analyzed by mass spectrometry. HPLC-ESI-MS/MS was performed as previously described^39^. In brief, MS/MS was performed in positive ion mode on a Thermo Scientific Orbitrap Fusion Lumos tribrid mass spectrometer fitted with an EASY-Spray Source (Thermo Scientific, San Jose, CA). NanoLC was performed using a Thermo Scientific UltiMate 3000 RSLCnano System with an EASY Spray C18 LC column (Thermo Scientific).

Tandem mass spectra were extracted from Xcalibur ‘RAW’ files and charge states were assigned using the ProteoWizard 2.1.x msConvert script using the default parameters(23). The fragment mass spectra were then searched against the *Saccharomyces cerevisiae* (strain ATCC 204508 / S288c) (Baker’s yeast) UniProt database (6067 entries) using Mascot (Matrix Science, London, UK; version 2.6) using the default probability cut-off score. Cross-correlation of Mascot search results with X! Tandem was accomplished with Scaffold (version Scaffold_4.8.7; Proteome Software, Portland, OR, USA). Probability assessment of peptide assignments and protein identifications were made through the use of Scaffold. Only peptides with ≥ 95% probability were considered. Progenesis QI for proteomics software (version 2.4, Nonlinear Dynamics Ltd., Newcastle upon Tyne, UK) was used to perform ion-intensity based label-free quantification similar to as previously described^39^. Principal component analysis and unbiased hierarchal clustering analysis (heat map) was performed in Perseus^40,41^. Gene ontology and KEGG pathway enrichment analysis was performed with DAVID^42^.

### Recombinant Mpro and proteolytic activity assays

SARS-CoV-2 M^pro^ mutants were generated with QuikChange® II Site-Directed Mutagenesis Kit from Agilent (Catalog #200524), using plasmid pE-SUMO-Mpro as the template. The plasmid produces tag-free Mpro protein with no extra residue at either N- or C-terminus upon removal of the SUMO tag by SUMO protease digestion^17^.

SARS-CoV-2 M^pro^ mutant proteins were expressed and purified as previously described^17,24^ with minor modifications. Plasmids were transformed into E. coli BL21(DE3) competent cells and bacterial cultures overexpressing the target proteins were grown in LB (Luria-Bertani) medium containing 50 µg/mL of kanamycin at 37 °C, and expression of the target protein was induced at an optical density (A600) of 0.6-0.8 by the addition of isopropyl β-d-1-thiogalactopyranoside (IPTG) to a final concentration of 0.5 mM. The cell culture was incubated at 18°C for 12-16 hrs. Bacterial cultures were harvested by centrifugation (8,000 ×g, 10 min, 4°C) and resuspended in lysis buffer containing 25 mM Tris (pH 8.0), 750 mM NaCl, mM DTT, 0.5 mg/mL lysozyme, 0.5 mM phenylmethylsulfonyl fluoride (PMSF) and 0.02 mg/mL DNase I. Bacterial cells were lysed by alternating sonication (30% amplitude, 1 s on/1 s off) and homogenization using a tissue grinder. The lysed cell suspension was clarified by centrifugation (18,000 ×g, 30min, 4°C) and the supernatant was incubated with Ni-NTA resin for over 2 hrs at 4°C on a rotator. The Ni-NTA resin was thoroughly washed with 20 mM imidazole in washing buffer containing 50mM Tris (pH 8.0), 150mM NaCl, 2 mM DTT, and SUMO-M^pro^ protein was eluted with elution buffer containing 50 to 300mM imidazole, 50mM Tris (pH 8.0), 150mM NaCl, 2mM DTT. Fractions containing SUMO-M^pro^ proteins greater than 90% homogeneity were pooled and subjected to dialysis (two times) against a buffer containing 50mM Tris (pH 8.0), 150mM NaCl, 2mM DTT and 10% glycerol. SUMO protease digestion was carried out at 30ºC for 1 hr to remove SUMO tag. Following digestion, SUMO Protease and SUMO tag were removed by Ni-NTA resin. The purified tag-free SARS-CoV-2 Mpro mutant proteins were fast frozen in liquid nitrogen and stored at -80 °C.

For measurement of *K*_*m*_*/V*_*max*_ of SARS-CoV-2 M^pro^ mutants, proteolytic reactions were carried out with optimized concentrations of the mutant proteins and a series of concentrations of FRET substrate ranging from 0 to 200 µM in 100μL of reaction buffer containing 20mM HEPES (pH 6.5), 120mM NaCl, 0.4mM EDTA, 4mM DTT, and 20% glycerol at 30°C in a BioTek Cytation 5 imaging reader (Agilent) with filters for excitation at 360/40 nm and emission at 460/40 nm. Reactions were monitored every 90s, and the initial velocity of the proteolytic activity was calculated by linear regression for the first 15min of the kinetic progress curves. The initial velocity was plotted against the FRET substrate concentrations using the classic Michaelis-Menten equation in Prism 8 software.

For IC_50_ measurements, optimized concentrations of the mutant proteins were incubated with series concentrations of GC-376, PF-00835231 or nirmatrelvir (PF-07321332) in 100μL of reaction buffer at 30°C for 15 min, and the reaction was initiated by adding 10μM FRET substrate. The reaction was monitored for 1 hr, and the initial velocity was calculated for the first 15min by linear regression. The IC_50_ was determined by plotting the initial velocity against various concentrations of the compounds using log (inhibitor) vs response-variable slope in Prism 8 software.

For *K*_*i*_ measurements, optimized concentrations of the mutant proteins were added to 20μM FRET substrate with various concentrations of GC-376, PF-00835231 or nirmatrelvir (PF-07321332) in 200μL of reaction buffer at 30°C to initiate the proteolytic reaction. The reaction was monitored for 2 hrs and the initial velocity was calculated for the first 90 min by linear regression. The *K*_*i*_ was calculated by plotting the initial velocity against various concentrations of the compounds using Morrison plot (tight binding) in Prism 8 software.

### M^pro^ crystallization and structure determination

SARS-CoV-2 M^pro^ E166N/R was diluted to 5 mg/mL in protein buffer (50 mM Tris pH 7.0, 150 mM NaCl, 4 mM DTT). Protein for complex determination was incubated overnight at 4 °C with 2mM GC376. No precipitation was observed after incubation, and centrifugation was not necessary. Apo and complex crystals were grown using 1.5 μL:1.5 μL (protein:well solution) hanging drops and a well solution of 0.1 M MgCl_2_, 20% PEG 3350, 10% 1,6-hexanediol, 0.1 M HEPES pH 7.5, and 0.1 M LiSO_4_. E166N/R crystals grew overnight at 20 °C. Crystals were cryoprotected using the well solution supplemented with 20% glycerol, and then flash-frozen in liquid nitrogen.

X-ray diffraction data (Table S3) were collected at the Southeast Regional Collaborative Access Team (SER-CAT) 22-BM beamline at the Advanced Photon Source (APS) in Argonne, IL, and processed with HKL2000 and CCP4. PHASER was used for molecular replacement using a previously solved SARS-CoV-2 M^pro^ structure (PDB ID: 7LYH) as a reference model. The CCP4 suite, (23) Coot, (24) and the PDB REDO server (pdb-redo.eu) (25) were used to complete the model building and refinement. The PyMOL Molecular Graphics System (Schrödinger, LLC) was used to generate all images.

## Acknowledgements

This work was supported by the National Institutes of Health (NIH) grants 1R15GM129766-01 to J.S.C. and AI158775 to J.W. Additional support was provided by funds from the Adeline E. Fracassi Trust (Gregory A. Pope) and the Litowitz Family Fund to J.S.C. J.S.C. thanks Drs. Stephen J. Kron and Sean Palacek for critical discussions related to this work. We are grateful to the staff and scientists at SER-CAT for assistance with X-ray diffraction data collection. SEC-CAT is supported by its member institutions, and equipment grants (S10_RR25528, S10_RR028976 and S10_OD027000) from the National Institutes of Health. Use of the Advanced Photon Source was supported by the U. S. Department of Energy, Office of Science, Office of Basic Energy Sciences, under Contract No. W-31-109-Eng-38. J.S.C. dedicates this work to the loving memory of Yin Y. Choy.

## Author Contributions

J.S.C, Y.C., J.W. conceived the project, analyzed, and interpreted data. J.O. performed all yeast experiments. X.Z. constructed and purified mutants. R.T.M. crystallized mutants. E.M.L. analyzed crystals, collected diffraction data and determined the structures with assistance from L.M.C.J., M.J.B. YH and HT performed protein expression, purification and enzyme assays. A.A.L. and P.L. performed mass spectrometry and analysis. J.S.C. wrote the manuscript with input from Y.C. and J.W.

## Data Accessibility

The X-ray crystal structures have been deposited into the Protein Data Bank with accession codes 8DDI (E166N Apo) and 8DDM (E166R GC376).

