## Supplemental Information for "A yeast-based system to study SARS-CoV-2 M^pro^ structure and to identify nirmatrelvir resistant mutations"

**This PDF file includes:**

Tables S2-S4

Figures S1-S4

Table S1. Plasmids used in this study

| Number | [1]Plasmid | Gene |
|---|---|---|
| JCB462 | pGBW-m4046203 | SARS-CoV-2 $M^{pro}$ ($3CL^{pro}$) |
| JCB464 | pGBW-m4046418 | SARS-CoV-2 N (Nucleocapsid) |
| JCB465 | pGBW-m4046249 | SARS-CoV-2 NSP12 (RNA-dependent RNAP) |
| JCB467 | pGBW-m4046574 | SARS-CoV-2 S (Spike protein) |
| JCB504 | pGBW-m4046277 | SARS-CoV-2 M (Membrane protein) |
| JCB505 | pGBW-m4046231 | SARS-CoV-2 NSP8 |
| JCB506 | pGBW-m4046455 | SARS-CoV-2 E (Envelope protein) |
| JCB507 | pGBW-m4046246 | SARS-CoV-2 NSP7 |
| JCB508 | pGBW-m4046415 | SARS-CoV-2 NSP13 (Helicase) |
| JCB509 | pGBW-m4046483 | SARS-CoV-2 NSP3 ($PL^{pro}$) |

[1] Plasmids were gifts from Ginkgo Bioworks & Benjie Chen and obtained from Addgene.

Table S2. Primer Sequences

| Mutant | F or R | Sequence |
|---|---|---|
| C145A | Forward Primer | 5'-GGTTCTGCTGGCTCCGTTGGTTTTA-3' |
| | Reverse Primer | 5'-GGAGCCAGCAGAACCATTCAAAAAA-3' |
| E166R | Forward Primer | 5'-TCACATGAGATTGCCTACAGGTGTTCACGC-3' |
| | Reverse Primer | 5'-GGCAATCTCATGTGATGCATGTAACAGAAACTG-3' |
| E166N | Forward Primer | 5'-TCACATGAACTTGCCTACAGGTGTTCACGC-3' |
| | Reverse Primer | 5'-GGCAAGTTCATGTGATGCATGTAACAGAAACTG-3' |
| E166D | Forward Primer | 5'-TCACATGGACTTGCCTACAGGTGTTCACGC-3' |
| | Reverse Primer | 5'-GGCAAGTCCATGTGATGCATGTAACAGAAACTG-3' |
| P132H | Forward Primer | 5'-TATGAGGCACAACTTCACTATTAAGGGTTCTTTTT-3' |
| | Reverse Primer | 5'-AAGTTGTGCCTCATAGCGCACTGGTATACACC-3' |
| N142A | Forward Primer | 5'-TTTTTTGGCTGGTTCTTGTGGCTCCGTT-3' |
| | Reverse Primer | 5'- GAACCAGCCAAAAAAGAACCCTTAATAGTGAAG-3' |

Table S3. X-ray Data Collection and Refinement Statistics
Data Collection

| Structure (PDB ID) | 8DDI | 8DDM |
|---|---|---|
| Mutation | E166N | E166R |
| Ligand | Apo | GC376 |
| Space Group | C2 | C2 |
| Cell Dimensions | | |
| *a*, *b*, *c* (Å) | 116.138 | 113.931 |
| | 53.928 | 53.401 |
| | 44.636 | 46.482 |
| α, β, γ (°) | 90 | 90 |
| | 101.27 | 102.12 |
| | 90 | 90 |
| Resolution (Å) | 50-2.80 | 50-2.78 |
| No. Reflections | 6641 | 6825 |
| $R_{merge}$ (%) | 11.6 | 9.8 |
| *I* / *óI* | 12.18 (2.33) | 13.22 (2.11) |
| Completeness (%) | 98.2 | 97.7 |
| Redundancy | 3.8 (1.9) | 4.3 (2.4) |
| **Refinement** | | |
| Resolution (Å) | 43.81-2.80 | 48.20-2.78 |
| $R_{work}/R_{free}$ (%) | 17.8/24.7 | 19.1/24.6 |
| **No. Heavy Atoms** | | |
| Protein | 2376 | 2385 |
| Ligand/Ion | 0 | 29 |
| Water | 22 | 20 |
| ***B*-Factors (Å$^2$)** | | |
| Protein | 49.03 | 49.38 |
| Ligand/Ion | 0 | 43.41 |
| Water | 33.4 | 34.49 |
| **Ramanchandran Plot** | | |
| Favored Region (%) | 97.4 | 97.0 |
| Allowed Region (%) | 2.3 | 2.6 |
| Outlier Region (%) | 0.3 | 0.3 |

* Values in parentheses represent highest resolution shells

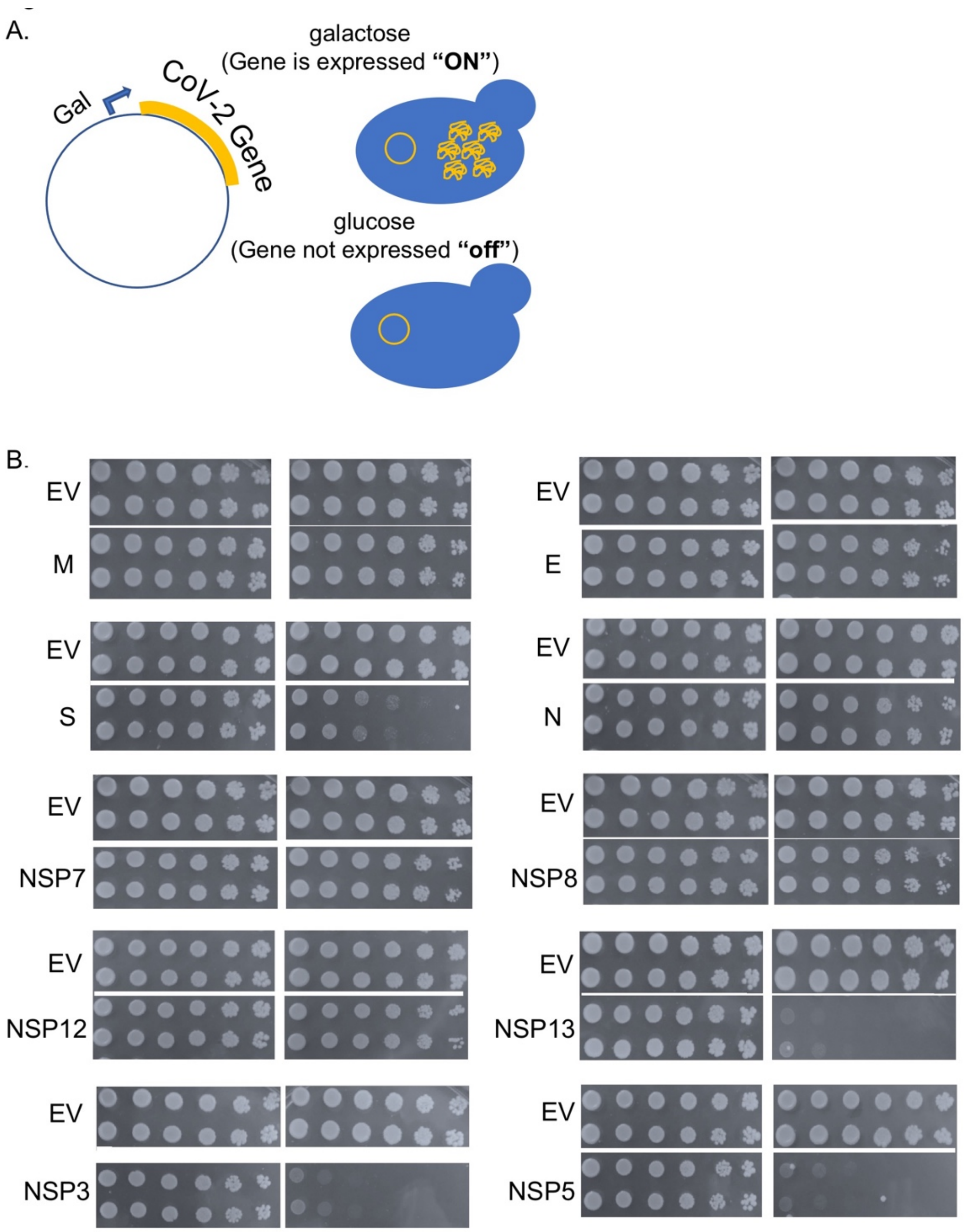

**Fig. S1. Expression of SARS-CoV-2 Genes in yeast.** A) SARS-CoV-2 genes were cloned into a high copy yeast plasmid and regulated by a galactose inducible promoter (Gal1). B) Spot assays were performed on glucose (left) or galactose (right) containing plates to determine the growth effects conferred by expression of the indicated genes. Two biological replicates were performed for each (two rows) EV and indicated gene. Samples (3ul) of a 5-fold serial dilution are spotted in each row. EV refers to yeast carrying an empty vector. M, E, S, N are structural proteins and all others are non-structural proteins (NSP).

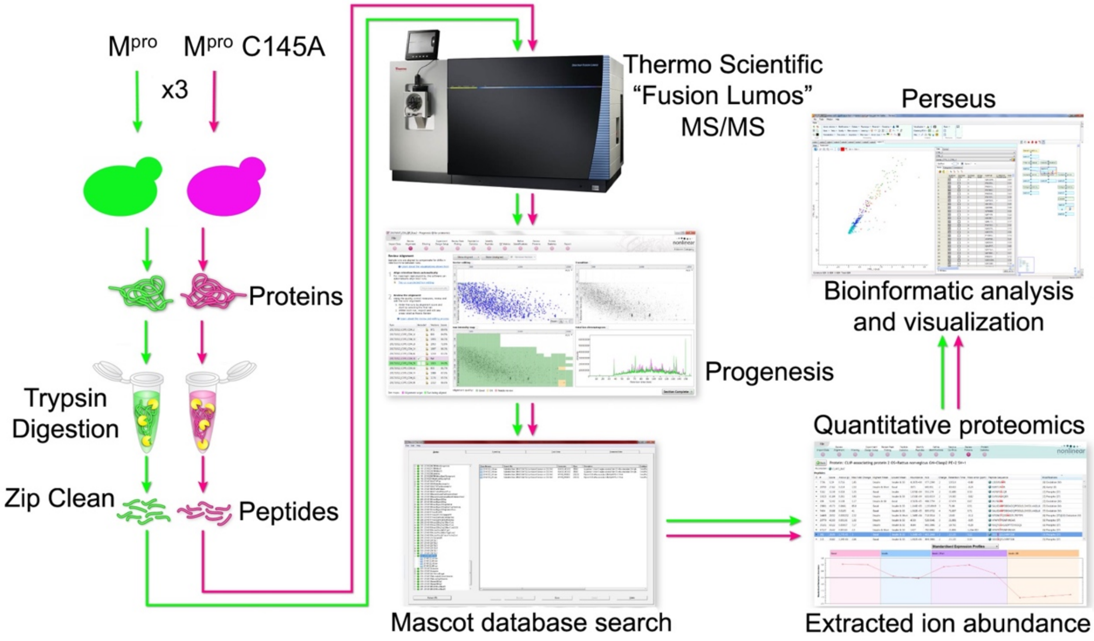

**Fig. S2. Determining potential yeast substrates of M$^{pro}$.** Schematic of the work flow to perform proteomics from yeast strains expressing wild-type M$^{pro}$ or catalytically inactive M$^{pro}$ C145A.

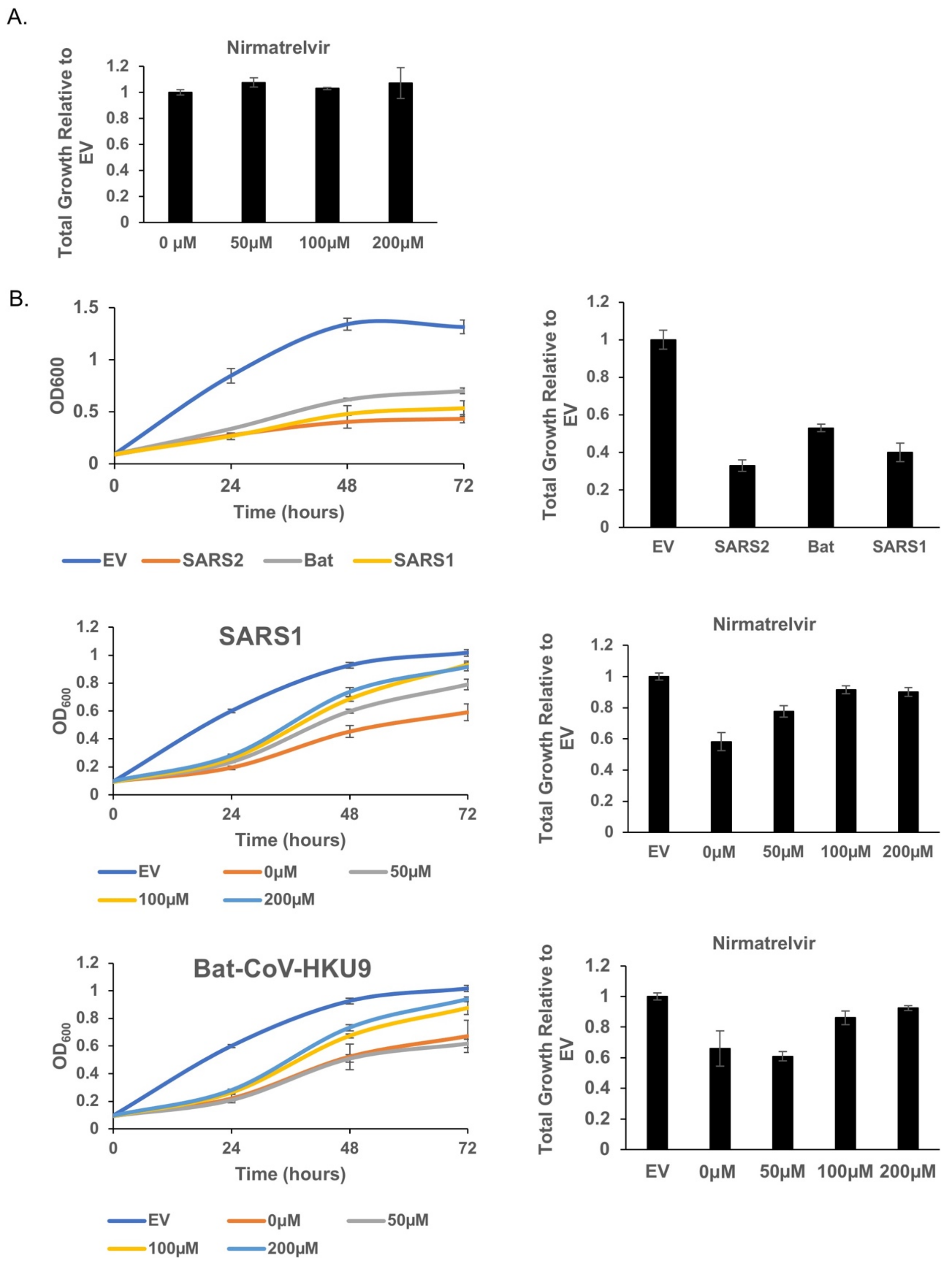

**Fig. S3. Effects of nirmatrelvir on yeast expressing Mpro from SARS-CoV-1 and Bat-CoV-HKU9.** A) No changes in growth were observed in yeast treated with nirmatrelvir up to 200μM. B) Growth of yeast expressing $M^{pro}$ from the indicated coronavirus is plotted in the presence or absence of nirmatrelvir. $M^{pro}$ from both viruses remain responsive to nirmatrelvir as indicated improved growth (nearly matching the empty vector (EV) control) in the presence of nirmatrelvir.

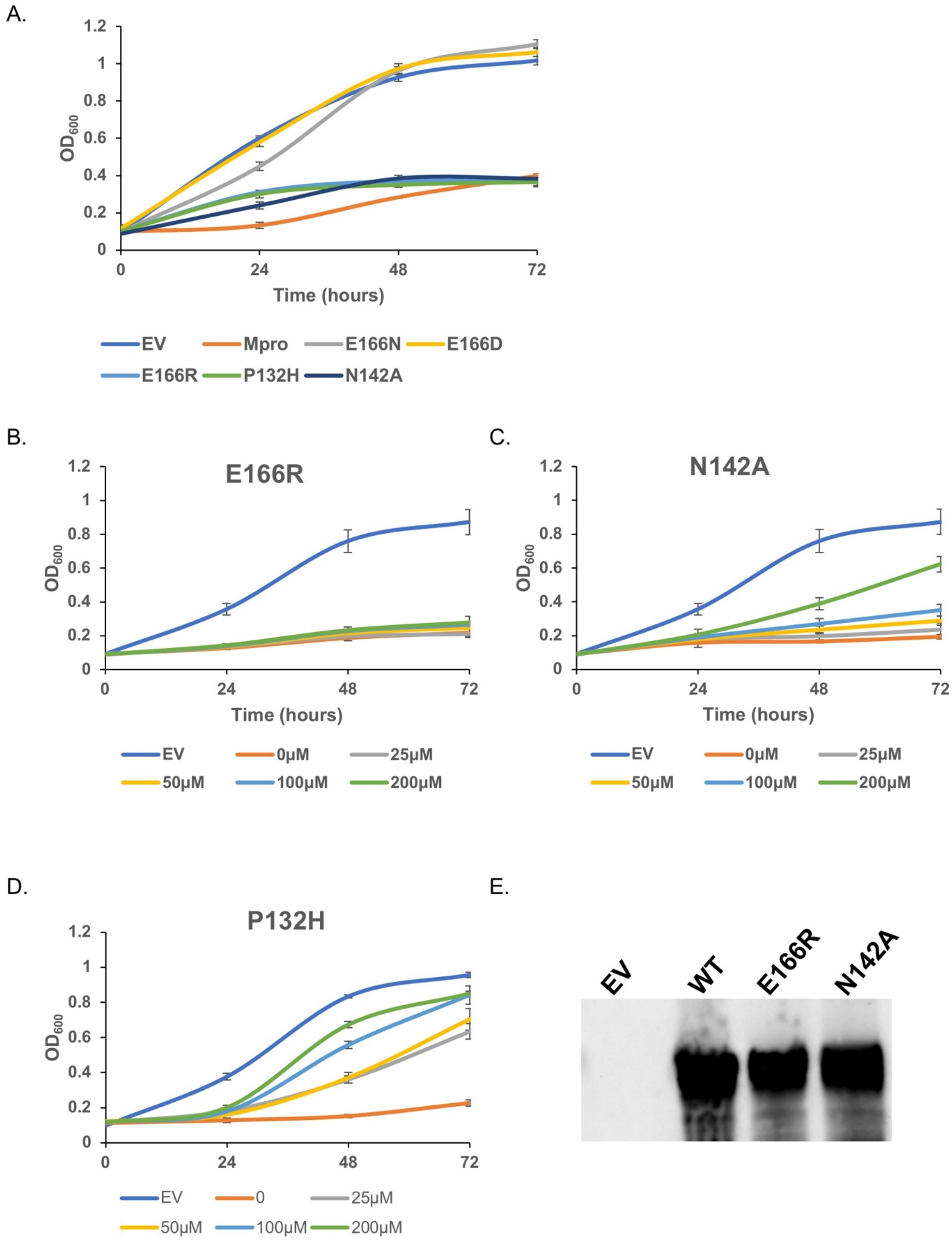

**Fig. S4. Growth curves for yeast expressing SARS-CoV-2 M^pro^.** A) Indicated M^pro^ mutants were expressed in yeast. Wild-type M^pro^, M^pro^ E166R, M^pro^ P132H, and M^pro^ N142A all confer a strong growth defect compared to empty vector (EV) control. On the other hand $M^{pro}$ E166N and $M^{pro}$ E166D do not confer a growth defect and grow as well as EV control. B-D) Growth curves for $M^{pro}$ E166R, $M^{pro}$ P132H, and $M^{pro}$ N142A treated with 0, 25, 50, 100, 200μM of nirmatrelvir. E) Western blot showing levels of wild-type Mpro, $M^{pro}$ E166R, and $M^{pro}$ N142A.
